# Harnessing population-specific protein truncating variants to improve the annotation of loss-of-function alleles

**DOI:** 10.1101/2020.08.17.254904

**Authors:** Rostislav K. Skitchenko, Julia S. Kornienko, Evgeniia M. Maksiutenko, Andrey S. Glotov, Alexander V. Predeus, Yury A. Barbitoff

## Abstract

Accurate annotation of putative loss-of-function (pLoF) variants is an important problem in human genomics and disease, which recently drew substantial attention. Since such variants in disease-related genes are under strong negative selection, their frequency across major ancestral groups is expected to be highly similar. In this study, we tested this assumption by systematically assessing the presence of highly population-specific protein-truncating variants (PTVs) in human genes using latest population-scale data. We discovered an unexpectedly high incidence of population-specific PTVs in all major ancestral groups. This does not conform to a recently proposed model, indicating either systemic differences in disease penetrance in different human populations, or a failure of current annotation criteria to accurately predict the loss-of-function potential of PTVs. We show that low-confidence pLoF variants are enriched in genes with non-uniform PTV count distribution, and developed a computational tool called LoFfeR that can efficiently predict tolerated pLoF variants. To evaluate the performance of LoFfeR, we use a set of known pathogenic and benign PTVs from the ClinVar database, and show that LoFfeR allows for a more accurate annotation of low-confidence pLoF variants compared to existing methods. Notably, only 4.4% of protein-truncating gnomAD SNPs in canonical transcripts can be filtered out using a recommended threshold value of the recently proposed *pext* score, while up to 10.9% of such variants are filtered using LoFfeR with the same false positive rate. Hence, we believe that LoFfeR can be used for additional filtering of low-confidence pLoF variants in population genomics and medical genetics studies.

## Introduction

International human genome resequencing projects have greatly enhanced our understanding of human genome variation. Rapid advances in sequencing technologies and data analysis led to the emergence of large-scale variant compendia, such as the Genome Aggregation Database (gnomAD) (Lek et al., 2016; Karczewski et al., 2019), that aggregate variation from many tens of thousands of individuals. These global variant datasets are successfully used by geneticists to gain novel insights into the evolutionary constraint of human genome. Diverse statistical models have been proposed to estimate the selective pressure for both whole genes (Lek et al., 2016; Cassa et al., 2017) and smaller genomic regions (Havrilla et al., 2018). Some of these methods use a specific class of genetic variants, namely the protein-truncating variants (PTVs), to evaluate evolutionary conservation of individual genes (Cassa et al., 2017). PTVs are highly deleterious genetic variants that frequently lead to the loss of gene product function (loss-of-function mutations, LoF). Thus, a large proportion of PTVs cause hereditary genetic conditions and are of ultimate relevance for variant interpretation in clinical diagnostics (Richards et al., 2015; Nykamp et al., 2017).

One other important type of information used in variant interpretation is the populational alternative allele frequency (AF). Despite that gnomAD-derived global AF estimates are frequently used for this purpose, it is widely anticipated that maximum AF among human subpopulations (popmax AF) should be taken into account as well (Lek et al., 2016). It is important to note, however, that while genetic drift can greatly affect allele frequencies in different ancestral groups for neutral or nearly neutral genetic variants, LoF variant counts should be distributed more uniformly due to an active selective pressure. Hence, the presence of highly population-specific LoF variants could indicate either (i) unequal evolutionary constraint in different populations, or (ii) variant misclassification, i.e. erroneous annotation of a variant as a genuine LoF. Both of these issues could directly translate into costly mistakes in clinical practice during variant interpretation. To this end, we set off to evaluate the presence of population-specific PTVs in gnomAD data, and suggest a method to avoid mistakes during clinical interpretation of such variants.

## Materials and methods

### Data acquisition and filtering criteria

We used the Genome Aggregation Database (gnomAD) release 2.1 exome data (accessed 2019-01-08) for the analysis. We have reimplemented the previously proposed framework for the selection of PTVs (Cassa et al., 2017). To this end, we selected single nucleotide substitutions annotated as nonsense (stop_gained) or splice site (splice_donor or splice_acceptor) mutations. Variants that were either flagged by LOFTEE (https://github.com/konradjk/loftee) or not annotated as PTVs in the canonical transcript were filtered. PTV counts in major gnomAD populations were aggregated for each gene after filtering. We then excluded all genes that have mean depth < 30x across gnomAD exomes, as well as genes with total PTV allele count (AC) > 0.001 □ allele number (AN). We then matched the resulting dataset with the ExAC-based data obtained by *Cassa et al*., omitting all genes not analyzed in either dataset.

### Statistical assessment of allele count distribution

Statistical analyses and modeling were done using R v. 3.6.1 and Python v. 3.6. Plots were generated using the ggplot2 package for R (Wickham, 2016).

We used previously proposed deterministic framework for the model-based analysis of PTV count distribution (Cassa et al., 2017). To estimate the selection coefficient against heterozygous protein-truncating variants (s_het_), we applied a Bayesian framework described by Cassa et al. Briefly, we first fitted an Inverse Gaussian (IG) distribution to the observed global PTV count data by maximizing the following likelihood:

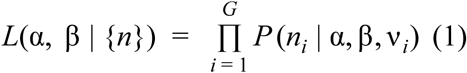

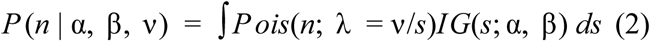

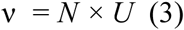

where *n* is the count of PTV alleles, *N* is the allele number (AN), U is the PTV-specific mutation rate (Samocha et al., 2014), and *s* is the selection coefficient against heterozygous PTV variants, *{n}* represents a set of PTV allele count across *G* genes in the dataset, and *α* and *β* are mean and shape parameters of the inverse Gaussian distribution, respectively.

Parameter fitting was done in each tercile of mutation rate distribution, as suggested in the original analysis. We then used the obtained IG parameters to infer the s_het_ value by maximizing the probability:

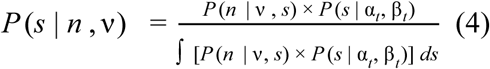

where *α*_*t*_ and *β*_*t*_. are the inferred IG parameters for a given mutation rate tercile.

To evaluate whether the observed PTV count distribution across major gnomAD populations fits the deterministic PTV count model, we calculated the likelihood of PTV count distribution (cross-population likelihood or CPL) by integrating the product probability of each s_het_ value given PTV allele counts in five major ancestral groups:

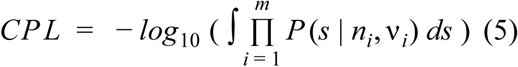

where *m* is the number of populations considered, *P(s* | *n*_*i*_, *v*_*i*_,) denotes the posterior probability of an *s*_*het*_ value given the observed and expected PTV count given by eq. (4).

Final observation likelihood score (OLS) was calculated as a *Z*-score of *CPL* value estimated from empirical *CPL* distribution parameters. These parameters were obtained by calculating *CPL* value for 100 random samples of PTV allele counts for each population *i* from Poisson distribution with mean *ν*_*i*_ */s*.

### Functional features of pLoF variants

To investigate functional properties of selected PTVs, we used the map of evolutionary constrained coding regions (CCR) (Havrilla *et al*., 2018), and the complete Genotype Tissue Expression (GTEx, v7) (Deluca et al., 2015) consortium expression dataset.

CCR percentile values were aggregated across all GENCODE v19 CDS regions to obtain estimates of evolutionary constraint for each exon separately. We then calculated the normalized per-exon constraint value by dividing mean exon CCR percentile by the maximum per-exon CCR percentile for each gene. Variants were then assigned to exons, and the most constrained exon was selected in case when two or more CDS regions overlapped a PTV.

For each transcript in the GTEx expression dataset, we first calculated mean and maximum expression levels in TPM for each isoform across all samples. We then selected (i) the isoform with the highest mean expression for a given gene; and (ii) the isoform with the highest maximum level of expression. pLoF variants were then annotated using binary variables indicating whether they affect these isoforms.

Base-level proportion expression across transcripts (*pext*) scores and transcript-level constraint measures were obtained from the gnomAD website: http://gnomad.broadinstitute.org/downloads/ (accessed 2020-02-12). For each PTV considered, we selected sets of isoforms that were either affected or unaffected by a pLoF variant. We then calculated the minimum LOEUF value and maximum pLI value across affected and unaffected isoforms for each variant.

### Model training and evaluation

Random forest was used as a default classification algorithm throughout the analysis. Prior to model fitting, a training set with balanced classes was constructed from the initial data using a downsampling procedure. Model performance was evaluated using ROC/AUC analysis with 3-fold to 10-fold cross validation. Receiver-operator (ROC) curves and area under curve (AUC) values were calculated using the pROC R package (Robin et al., 2011).

To create a validation dataset based on ClinVar variants, we selected protein-truncating variants from ClinVar (release 2020-06-15) using the same criteria as for the selection of PTVs from the gnomAD dataset. Prior to variant selection, a VCF file with ClinVar variants was annotated using Ensembl VEP v. 98.3 with the following plugins: LOFTEE, MPC, CADD, ExACpLI. Variants that were reported as pathogenic by multiple submitters, as well as known benign variants were included into the dataset for model training and evaluation. Only variants without conflicting interpretations were used.

When assessing the false-positive rate for different filtering approaches, we calculated the number of filtered pathogenic variants in the ClinVar dataset, and divided this number by the total number of pathogenic variants in the dataset (3,483). All variants for which the prediction could not be made were considered as high-confidence pLoF.

### Data and code availability

All code pertinent to the data analysis presented can be found at http://github.com/mrbarbitoff/ptv_project_bi/.

## Results

To evaluate the presence of population-specific PTVs (psPTVs) in gnomAD, we first selected all protein-truncating variant sites according to the criteria proposed by Cassa et al. (see Materials and Methods). Our initial selection resulted in 198,203 PTV sites belonging to 19,476 genes; after all filtering steps (see Materials and Methods), 140,287 PTVs and 15,844 genes were retained in the final dataset. We observed high similarity in the form of per-gene PTV allele count distributions in gnomAD subpopulations (Figure 1a) (except Finnish (FIN) and Ashkenazi Jewish (ASJ)), with the median of the distribution being roughly proportional to the sample size (allele number, AN).

**Figure 1.**
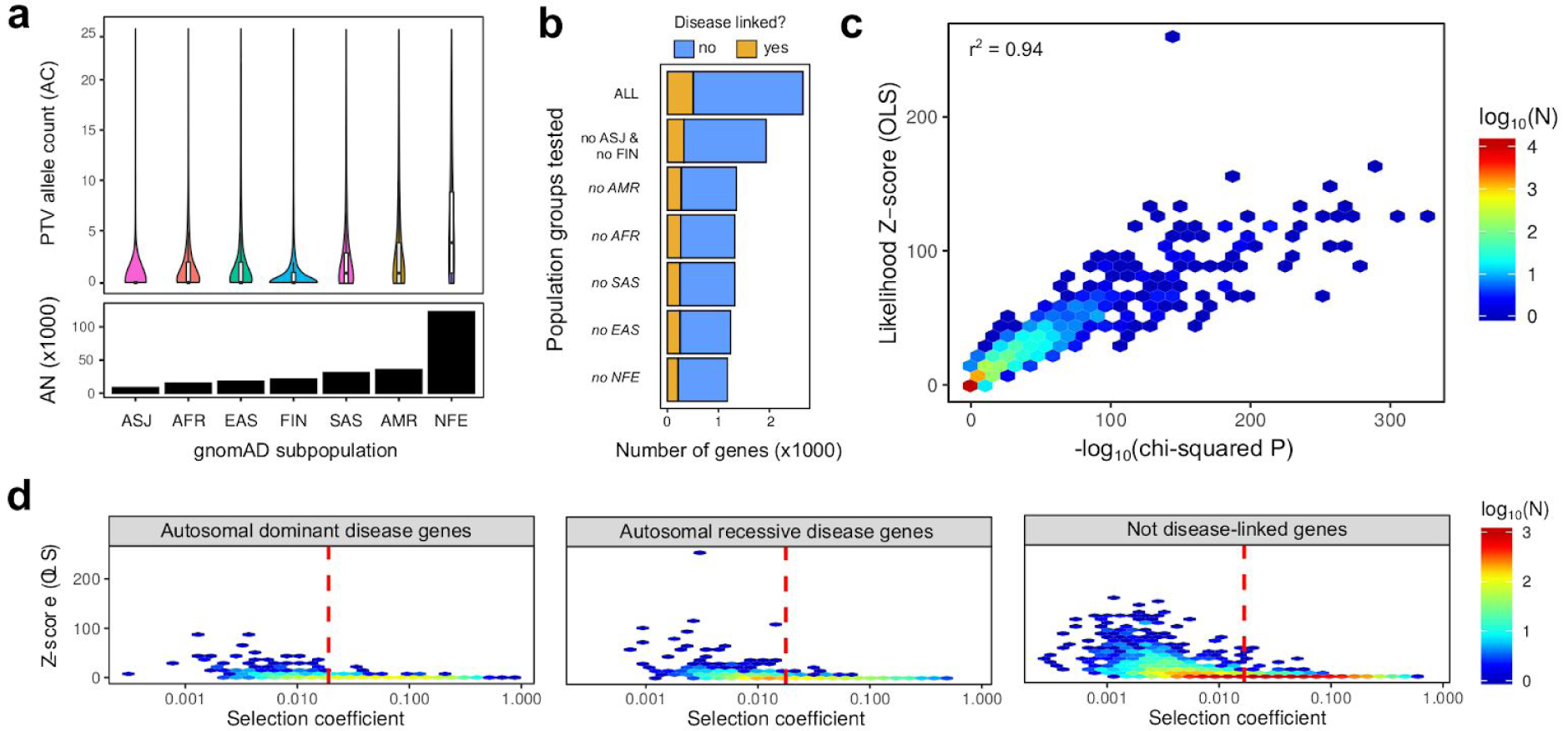
Identification and characterization of population-specific protein-truncating variant (psPTV) sites. **(a)** Distribution of per-gene PTV allele counts (AC) and mean allele numbers (AN) in major gnomAD populations. **(b)** Number of genes with significantly non-uniform distribution of PTV counts across gnomAD populations in the respective comparisons. Italic font indicates that testing was done in the dataset without FIN and ASJ populations. **(c)** Hexagonal scatterplot showing the correspondence between -log_10_(chi-squared p-value (test without FIN and ASJ) and the observation likelihood scores (OLS) for each gene. Fill is proportional to the log_10_ of gene count. **(d)** Hexagonal scatterplot showing the relationship between OLS and s_het_ values for genes associated with autosomal dominant (AD) or autosomal recessive (AR) disorders, as well as not associated with any disorder (according to OMIM). Dashed line corresponds to a threshold of s_het_ = 0.02, representing a sufficiently high value that allows to completely neglect the effects of genetic drift (Weghorn et al., 2019).

We then evaluated the uniformity of PTV count distribution across populations. To this end, we applied a chi-squared test to compare AC and AN values. Chi-squared test identified 2,710 genes showing significant (p < 1 * 10^−5^) non-uniformity of PTV allele counts (Figure 1b). Of these, 518 genes had known monogenic disease associations according to OMIM. Given these results, we questioned whether this non-uniformity is driven by the relatively inbred FIN and ASJ subgroups. Removal of these populations from the analysis decreased the number of genes with non-uniform PTV counts by a quarter; however, 1,972 genes still showed substantial deviations from uniformity (333 genes with known disease associations). We then asked if any population other than FIN and ASJ could account for this observation. Removing each of the five major ancestral groups (African (AFR), Ad Mixed American (AMR), East Asian (EAS), South Asian (SAS), and non-Finnish European (NFE)) decreased the number of genes with psPTVs (Figure 1b), but did not eliminate such genes. We thus concluded that psPTV sites are present in all major ancestral groups.

We then set off to evaluate whether genes with psPTV sites conform to the recently proposed model (Cassa et al., 2017). To this end, we first estimated the strength of selection against heterozygous protein truncating variants (s_het_) in each gene (see Materials and Methods). We observed modest differences in the inferred s_het_ distribution parameters (Figure S1) and the estimated s_het_ values (Figure S2 and Figure S3) between gnomAD and ExAC dataset, with the gnomAD-derived s_het_ distribution having higher median s_het_ and lower variance of estimated values. We then used estimated s_het_ values for each gene to calculate the likelihood of observed PTV count distribution across five major populations (see Materials and Methods for more details of the method). We further compared the estimated likelihoods with 100 random simulation replicates (see Methods). Our simulations showed that for most genes the correspondence between observed and expected likelihood values was high (Figure S4). We observed a high degree of correlation between chi-squared p-values and the Z-score of observed likelihood (observation likelihood score, OLS; r^2^ = 0.94) (Figure 1c), indicating that genes with high load of psPTVs do not conform to the established evolutionary model.

Deviation from the PTV evolutionary model can be explained by several major factors. One such factor is the decreased strength of natural selection which increases the effects of genetic drift on observed PTV counts (Charlesworth and Hill, 2018). To evaluate the influence of this factor on the observed non-uniformity of PTV counts, we analyzed the relationship between OLS and s_het_ values. Our analysis showed that, while many of the genes with high OLS either have low s_het_ or have no known disease association (Figure 1d), a substantial fraction of AD- and AR-disease associated genes under strong selection (s_het_ > 0.02) have high burden of psPTVs. As thus, violation of the PTV evolutionary model is unlikely to be explained solely by genetic drift.

The other possible explanation of the evolutionary model violation is the misannotation of PTV as a loss-of-function mutation. We first assessed whether any existing method could confirm misannotation of variants as genuine LoF ones. To this end, we applied the Annotation of Loss-of-function Transcripts (ALoFT) (Balasubramanian *et al*., 2017), one of the most recent computational toolkits for annotation of LoF variants. Surprisingly, we found that ALoFT not only did not show a higher fraction of tolerated pLoF variants in “violator” genes (OLS > 5) but, in contrast, showed a higher probability of dominant LoF effect for variants in such genes (Figure S5).

Given this unusual observation, we sought functional evidence that would indicate a higher proportion of low-confidence LoF variants in genes that show significant deviation from the model. We hypothesized that alternative splicing might play an important role in the observed effects. To check this hypothesis, we first compared the number of transcriptional isoforms and exons for violator and non-violator genes. As expected, we discovered that violator genes had more transcriptional isoforms and higher per-gene exon count compared to non-violator genes (Figure S6).

As psPTVs tend to localize in genes with higher number of isoforms, we also suggested that many of these variants might fall into transcriptional isoforms with low evolutionary constraint and/or expression levels. To test this hypothesis, we first compared the distribution of normalized per-exon constraint (CCR percentile) for variants located in genes with or without psPTV load. Indeed, we observed that PTVs in model-violating genes tend to reside in coding regions with lower evolutionary constraint (Figure 2a, leftmost panel), and the proportion of PTVs falling inside the most constrained exon in these genes is lower (Figure S7). We then compared the fraction of selected PTVs that fall into canonical isoforms with the highest average or maximum expression across GTEx samples (see Methods). We found that the fraction of variants that are found in such most expressed isoform is lower for genes harboring psPTVs (Figure 2a, second panel; Figure S7c).

**Figure 2.**
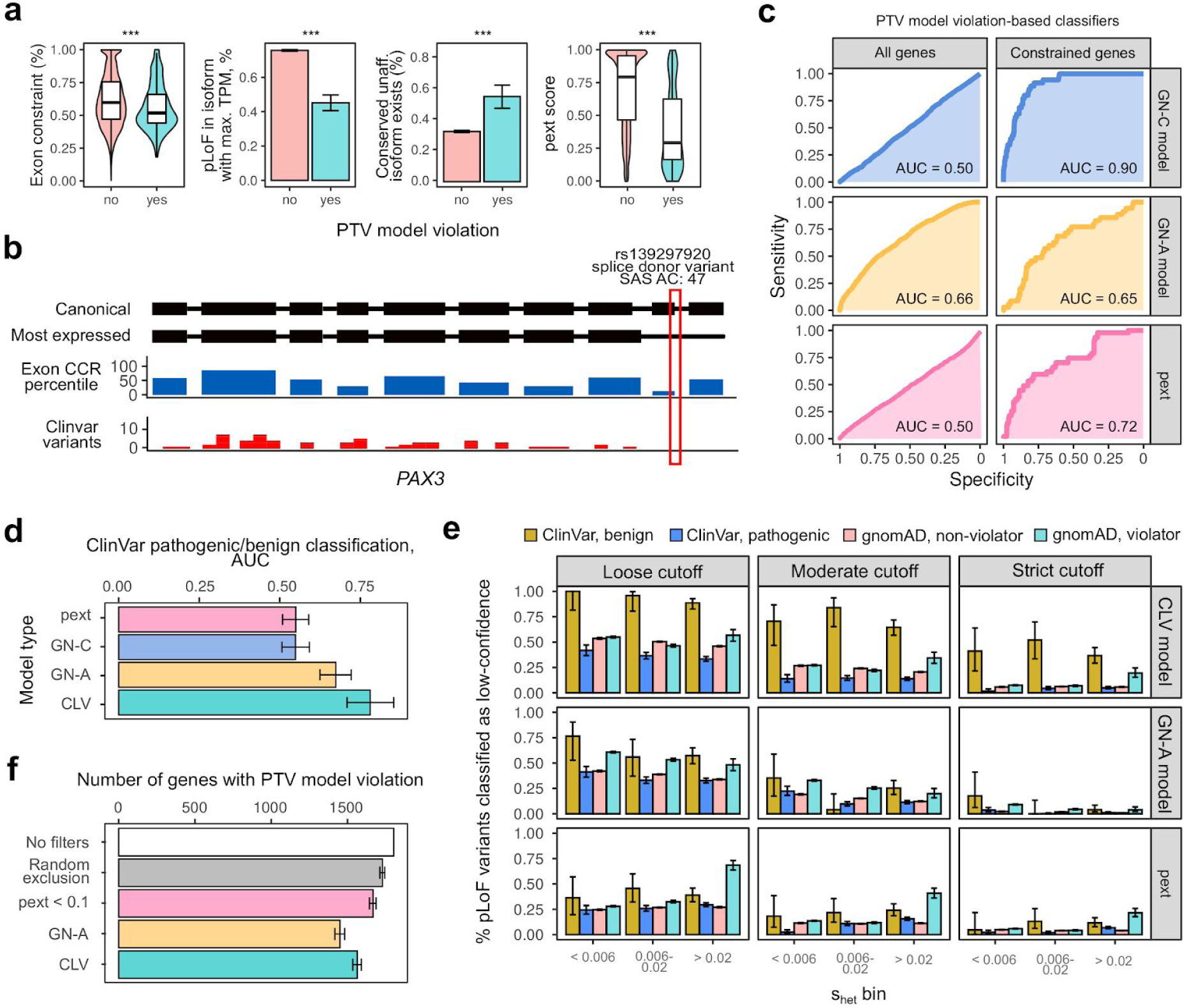
Construction of a machine learning model to predict low-confidence LoF variants based on PTV model violation. **(a)** Functional evidence of LoF misannotation for psPTVs. From left to right: violin plots representing normalized per-exon constraint (CCR percentile) value; fraction of PTVs falling to the most constrained exon; fraction of PTVs corresponding to a highly expressed transcript; fraction of PTVs corresponding to the transcript with the highest expression. Error bars represent 95% confidence interval for the binomial proportion. ***, p < 0.001. **(b)**. Schematic representation of a prevalent psPTV in the *PAX3* gene. **(c)** Receiver/operator curves (ROC) representing the performance of different approaches to classify variants in violator and non-violator genes. GN-C, a random forest model built using a training set of variants in constrained genes (s_het_ > 0.02); GN-A - same as GN-C, but trained using variants from all genes; pext - pext score cutoffs. Left part shows ROC curves obtained when evaluating performance using all gnomAD variants; right part shows ROC curves obtained using variants in highly constrained (s_het_ > 0.02) genes. **(d)** Area under curve (AUC) values from the ROC analysis of the classifiers’ performance on the validation dataset of the ClinVar variants. The CLV model is trained using a training set of variants derived from ClinVar itself. **(e)** Proportion of variants from ClinVar or gnomAD marked as low-confidence by different models. Error bars correspond to the 95% confidence intervals for binomial proportion. **(f)** Numbers of genes showing significant non-uniformity of PTV count distribution (chi-squared p-value < 1 * 10^−5^) before or after filtering of variants using indicated models. Filtering cutoffs were adjusted to exclude similar numbers of variants (40,000) in each case.

We next hypothesized that, given our observations, differences in functional properties of pLoF variants observed when comparing violator and non-violator genes with s_het_ > 0.02 should be less pronounced in genes with lower evolutionary constraint. Indeed, we found very little or no difference in the exon-level constraint and the proportion of variants affecting most expressed transcripts when comparing violator and non-violator genes with mild or no s_het_ filtering (Figure S7a-b). These results indicate that misannotation of LoF variants drives the deviation from the evolutionary model of PTV inheritance in highly conserved genes, while genetic drift seems to be more important in genes with lower conservation level. We also checked whether the observed differences between variants in violator and non-violator genes can be explained by one common psPTV. To do so, we analyzed whether removal of the most common PTV in each gene would eliminate the differences in exon-level CCR and expression of affected isoforms between variants in violator and non-violator genes. We found that the removal of the most common PTV did not have any effect on these functional properties (Figure S8), suggesting that the difference is not explained by one common population-specific variant site.

We then went on to compare major per-transcript constraint measures provided by gnomAD (pLI and LOEUF) for affected isoforms of violator and non-violator genes. We assumed that psPTVs not affecting the most expressed isoform might also affect less conserved transcriptional isoforms. To evaluate this hypothesis, we analyzed the maximum constraint values (i.e., minimum LOEUF and maximum pLI value) of transcripts affected by each PTV; and compared these values for violator and non-violator genes. Surprisingly, we found that PTVs in violator genes tend to affect isoforms with higher levels of evolutionary constraint (Figure S9). We then hypothesized that such unexpected result might be explained by the fact that the presence of common psPTVs in a gene might bias the estimates of s_het_, but not pLI and LOEUF, as s_het_ is based on total allele count of pLoF variants rather than on the number of pLoF. We therefore compared the relationship between s_het_ and pLi or LOEUF for violator and non-violator genes. Interestingly, we observed that violator genes tend to have lower s_het_ values than non-violator ones with similar pLI and/or LOEUF values (Figure S10), confirming our hypothesis.

We then questioned whether a highly conserved isoform unaffected by selected PTVs can be found for variants in violator and non-violator genes. To answer this question, we calculated the percentage of variants for which an unaffected isoform with LOEUF < 0.75 can be found; and compared this percentage for violator and non-violator genes. Indeed, we found that such conserved unaffected isoform can be found for a larger fraction of variants in violator genes as compared to non-violator ones (Figure 2a, third panel).

We also asked whether a recently proposed *pext* score that also represents the expression levels of isoforms affected by a PTV can explain the difference between variants in violator and non-violator genes. We observed a sharp difference in tissue-average pext scores between these gene groups, further supporting the assumption that LoF misannotation plays an important role in driving violations of PTV inheritance model in gnomAD (Figure 2a, rightmost panel).

Taken together, these results indicate that a substantial fraction of psPTVs are likely to be incorrectly annotated as LoF variants. One example of a psPTV that shows evidence for misannotation is the rs139297920 variant in the *PAX3* gene (Figure 2b). This variant is annotated as a donor splice site variant in the canonical transcript; however, this same variant falls into the 3’-UTR of the most expressed and most conserved isoform of *PAX3* (ENST00000350526), and corresponds to the exon with the lowest CCR percentile with no reported ClinVar variants.

As population-specificity of the PTVs can indicate their low LoF potential, we decided to build a predictive model to identify low-confidence LoF variants based on our data. We selected five major features to be used for predictive model construction: (i) normalized exon constraint level (Figure 2a, leftmost panel); (b) a binary variable indicating whether a PTV affects the isoform with the highest average expression; (Figure 2a, second panel); (c) a binary variable indicating whether a PTV affects the isoform with the highest maximum expression; (d) a binary variable indicating whether there is at least one unaffected isoform that has the same or higher LOEUF compared to affected ones; and (e) tissue-average pext score (Figure 2a, rightmost panel). We then constructed two random forest classifiers using either genes with s_het_ > 0.02 (gnomAD-based constrained model, denoted GN-C) or all genes in the dataset (all-gnomAD model, denoted GN-A). Next, we evaluated how well these classifiers discriminate between variants in violator and non-violator genes when considering a test set of gnomAD variants in all genes (Figure 2c, left) or in highly constrained (s_het_ > 0.02) genes (Figure 2c, right). We found that the GN-C model has the highest ROC/AUC value in constrained gene set (AUC = 0.90), while the performance of this classifier was very poor when using a test set derived from all genes. Similarly to the GN-C model, pext-only based classification also showed reasonable predictive power when classifying variants in constrained genes (AUC = 0.72) but performed much worse on the complete dataset (AUC = 0.51). In contrast to the GN-C and pext-only model, the GN-A classifier was able to discriminate between variants in violator and non-violator genes in both the complete dataset and for genes with high s_het_ (Figure 2c, AUC = 0.65 in both cases). Overall, we concluded that both GN-C and GN-A classifiers had some advantage over the pext-only model, with the GN-C model performing better on constrained genes, and the GN-A model - on all genes.

We next sought to prove that our classifiers indeed allowed for more efficient identification of low-confidence pLoF variants. To this end, we constructed a validation set of known pathogenic and benign ClinVar PTVs (see Methods). We then applied the constructed classifiers to this dataset to see whether our models would efficiently mark benign ClinVar PTVs as low-confidence LoF alleles. Both pext-only and the GN-C models were inefficient in separating benign and pathogenic pLoF variants (Figure 2d, Figure S11). At the same time, the GN-A model showed significantly better performance (AUC = 0.67), indicating that this model captures important information about low-confidence pLoF variants when trained to discriminate gnomAD variants in all violator and non-violator genes. We then asked if a more efficient random forest classifier with the exact same predictor variables can be constructed using a subset of the ClinVar data during model training. Indeed, we found that such a classifier (denoted as CLV) performed notably better compared to the GN-A model (AUC = 0.75). Since the CLV model was less efficient at classifying gnomAD variants (AUC = 0.52), we concluded that both GN-A and CLV models can be useful for more efficient annotation of low-confidence pLoF variants.

We then went on to assess the proportion of gnomAD and ClinVar PTVs marked as low-confidence pLoF by our models with varying prediction cutoffs for genes with different constraint levels. As expected, the majority of benign ClinVar variants in all three gene groups were marked as low-confidence LoF variants by the CLV and the GN-A models at the most permissive cutoff (prediction probability > 0.25), but not when filtering with pext < 0.5 which could only discriminate between pathogenic and benign variants in genes with moderate s_het_ values (0.006 to 0.02) (Figure 2e). At the same time both the CLV model and pext-based filtering could efficiently differentiate variants in violator and non-violator genes only for genes with the highest constraint values (s_het_ > 0.02), in contrast to the GN-A model which was efficiently separating such variants in all gene groups.

Cummings et al. have proposed a tissue-average pext value of 0.1 as a cutoff to be used for routine variant filtering. To evaluate the potential of our models for a similar purpose, we sought to identify threshold values of prediction probability for both GN-A and CLV models that would have similar false-positive rate as filtering by pext < 0.1. Around 4.9% of known pathogenic ClinVar variants were filtered out when using a pext threshold of 0.1. Such a false positive rate corresponds to the prediction probability cutoff of 0.575 for the GN-A and 0.59 for the CLV model. Applying these threshold values to filter out variants from the gnomAD dataset we found that 10.9% (15,309) of all gnomAD variants were filtered out using the GN-A model, and around 9.8% (13,695) with the CLV model. These values are 2.2 - 2.5 times higher than ones obtained when filtering with pext < 0.1 (6,168 variants; 4.4% of all gnomAD variants).

Finally, we asked whether our filtering strategy could substantially decrease the number of genes harboring psPTVs. To answer this question, we applied four different filtering approaches to the gnomAD dataset: (i) filtering by pext < 0.1; (ii) filtering with a matched cutoff value of the GN-A prediction probability; (iii) filtering with a matched cutoff value of the CLV prediction probability; and (iv) random exclusion of 6,168 variants from the data. Indeed, all filtering strategies could decrease the total PTV allele count for a large number of genes; notably, random exclusion tended to change the total allele count proportionally, while filtering with the other approaches resulted in more profound non-proportional changes (Figure S12). All approaches decreased the number of genes with non-uniform PTV count distribution as judged by chi-squared p-value (Figure 2f). Importantly, pext-based filtering performed only slightly better than random exclusion of variants (Figure 2f), while GN-A and CLV-based filtering allowed for more efficient elimination of the imbalance in PTV counts across populations. Moreover, we observed a small but significant correlation between the prediction probabilities for the GN-A and CLV models and the variant allele count (r^2^ = 0.053 for the GN-A model and 0.026 for the CLV model) (Figure S13), indicating that our models tend to mark PTVs with higher allele frequency as low-confidence LoF alleles without any prior knowledge of allele frequency.

To predict low-confidence pLoF variants in any custom VCF file, we implemented a simple bioinformatic tool called LoFfeR that applies both the GN-A and the CLV models to the data and classifies the variants according to the aforementioned classification cutoffs. The tool is freely available at https://github.com/mrbarbitoff/ptv_project_bi/.

## Discussion

The importance of population-level genetic data for the interpretation of variants in both research and clinical practice has been noted multiple times (e.g. in Lek *et al*., 2016). Population-specific allele frequency data might be important when classifying missense or other types of variants with disease-causing potential. At the same time, loss-of-function (LoF) variants in functionally important genes should be uniformly distributed across major populations (Weghorn *et al*., 2019). Nevertheless, we discovered prevalent psPTVs for 1,972 genes in the gnomAD dataset.

Highly population-specific LoF variants in a certain gene might arise due to several major factors, namely (i) genetic drift; (ii) difference in selection coefficients across populations; and (iii) misannotation of PTVs as LoF mutations. Genetic drift is a major factor contributing to derived allele frequency, and the possible effects of drift on the variance of observed PTV counts has already been discussed (Charlesworth and Hill, 2018). However, recent analysis indicates that including the effects of genetic drift into the model does not substantially affect s_het_ inference and PTV count variance for genes under strong selection (Cassa et al., 2018; Weghorn et al., 2019). As such, deviation from the deterministic model of PTV inheritance for genes with high s_het_ values indicates either decreased selection against LoF variants in a certain population or misannotation of LoF effect for a variant or a group of variants.

Importantly, the overall variance in selection coefficients and PTV allele counts decreases from ExAC to gnomAD (Figures S1 - S3), suggesting that increasing the sample size allows for more accurate inference of selective pressure acting on protein-coding genes. This observation implies that additional data aggregation efforts (like gnomAD v.3 and upcoming versions) should dramatically increase our ability to classify and interpret genetic variation.

Several lines of functional evidence indicate that, at least in part, psPTVs comprise variants that do not lead to a complete loss of gene function. Firstly, genes with high burden of psPTVs tend to have more transcriptional isoforms and exons (Figure S6). Secondly, PTVs in such genes tend to localize in exons with lower evolutionary constraint and not to disrupt at least one conserved isoform (Figure 2a). Thirdly, these variants tend not to affect the most expressed isoform and have lower *pext* scores (Figure 2a). Overall, we can conclude that an unusually high frequency of a PTV in a particular population (especially in genes with high overall constraint (i.e., high pLI and/or low LOEUF)) suggests that this variant is probably not a LoF allele.

Several methods have recently been proposed to identify and filter such low-confidence LoF alleles. These methods include the ALoFT software (Balasubramanian et al., 2017) and *pext* score (Cummings et al., 2020). These methods are of great interest to both medical and population genetics studies; however, neither ALoFT nor pext are sufficient for filtering of low-confidence LoF variants, as shown in our analysis (Figure 2c, Figure 2f, Figure S5). Moreover, the *pext* score has very low power to discriminate between known pathogenic and benign PTVs from ClinVar (Figure 2d). In this work we presented LoFfeR, a tool that utilizes two complementary random forest classifiers to enhance the prediction of genuine LoF variants. We showed that both models implemented in LoFfeR are capable of efficiently prioritizing low-confidence and benign pLoF variants (Figure 2c-d) and, hence, can be applied for routine variant filtration in clinical and research practice. Moreover, LoFfeR is substantially more efficient in eliminating the population-specific protein-truncating variants compared to *pext* (Figure 2f).

Notable examples of low-confidence pLoF variants filtered by LoFfeR (but not by *pext* alone) include the rs769874876 variant in the *PAX6* gene. This variant is present as a singleton PTV in gnomAD; however, similarly to the previously described rs139297920 in *PAX3*, rs769874876 does not affect the most expressed transcript and falls within the exon with no reported ClinVar pathogenic variants (while 165 pathogenic and likely pathogenic variants reported in the rest of the gene). Another interesting example is the SAS-specific rs570512700 splice site variant in the *TERF1* gene encoding a telomerase repeat binding factor 1. The variant occurs 74 times across gnomAD exomes, despite the fact that only 2 other pLoF sites are present in this gene (LOEUF = 0.37). The rs570512700 variant affects a highly expressed canonical isoform of *TERF1*, but does not affect the isoform with the highest mean expression (average pext = 0.41). Notably, exclusion of this very common variant results in a sharp increase in the estimated s_het_ value (0.0412 compared to 0.0029 before exclusion). This observation highlights the fact that the presence of population-specific and low-confidence LoF variants hinders accurate estimation of evolutionary constraint for human genes.

A substantial fraction of psPTVs can be filtered out with LoFfeR; however, for a large percentage of psPTVs, no clear explanation of their high frequency can be found. This observation, in turn, might be explained by the fact that the relationship between PTVs and loss of protein function is not explicitly understood. It is obvious that not all PTVs completely abrogate protein functionality (notable examples can be found in deep mutational scans of *TP53* (Kotler et al., 2018) and *TPMT* (Chiasson et al., 2019)). Various methods (such as MutPred-LoF (Pagel et al., 2017)) are being developed to predict the functional effect of a nonsense or frameshift variant on the functionality on the protein level. However, lack of large functional variant effect datasets complicates the task of predicting LoF effects of PTVs. The development of sophisticated experimental systems, such as the Variant Effect Mapping (VEM) in yeast (Welle et al., 2017), and construction of databases of such experimental functional evidence (Esposito et al., 2019), should greatly enhance our understanding of loss-of-function genetic effects. Integration of protein-based and transcript-/sequence-based methods for annotation of loss-of-function alleles might further improve the accuracy of variant interpretation in human medical and population genetics research.

## Acknowledgements

This study was supported by D.O. Ott Research Institute of Obstetrics, Gynaecology, and Reproductology, project 558-2019-0012 of FSBSI and by the Systems Biology Fellowship to Y.A.B.

