## Supplementary Figures for "Harnessing population-specific protein truncating variants to improve the annotation of loss-of-function alleles"

for

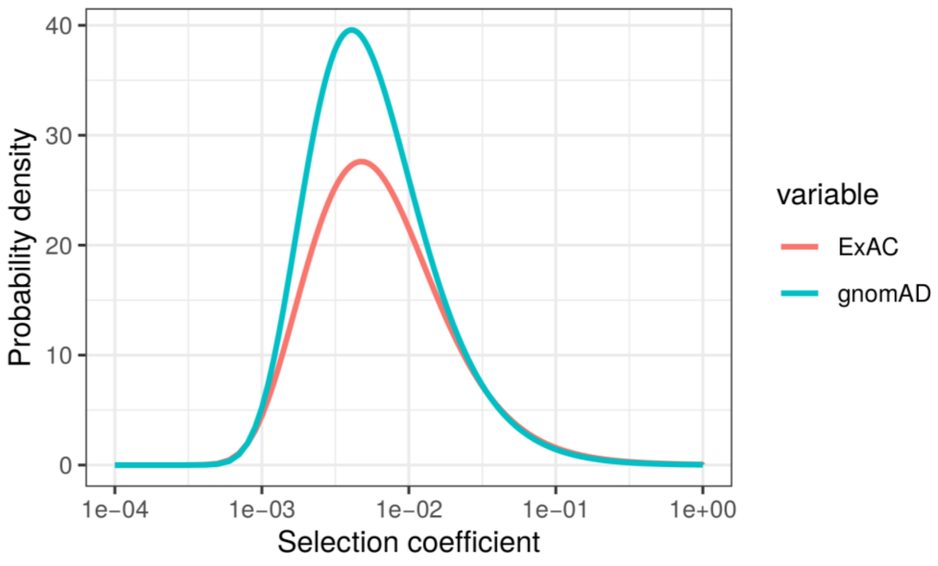


**Figure S1.** Distributions of selection coefficient ($s_{het}$) values estimated from ExAC data by Cassa et al., 2017 (red curve); and from gnomAD r.2.1 data in the present study (blue curve).

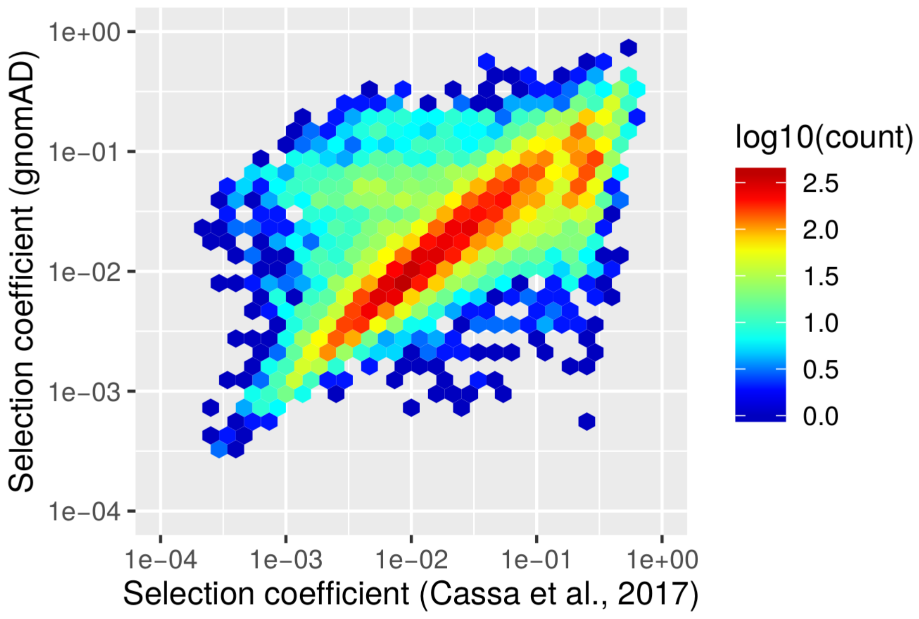


**Figure S2.** The correspondence between ExAC-based and gnomAD-based $s_{het}$ values estimated in this study.

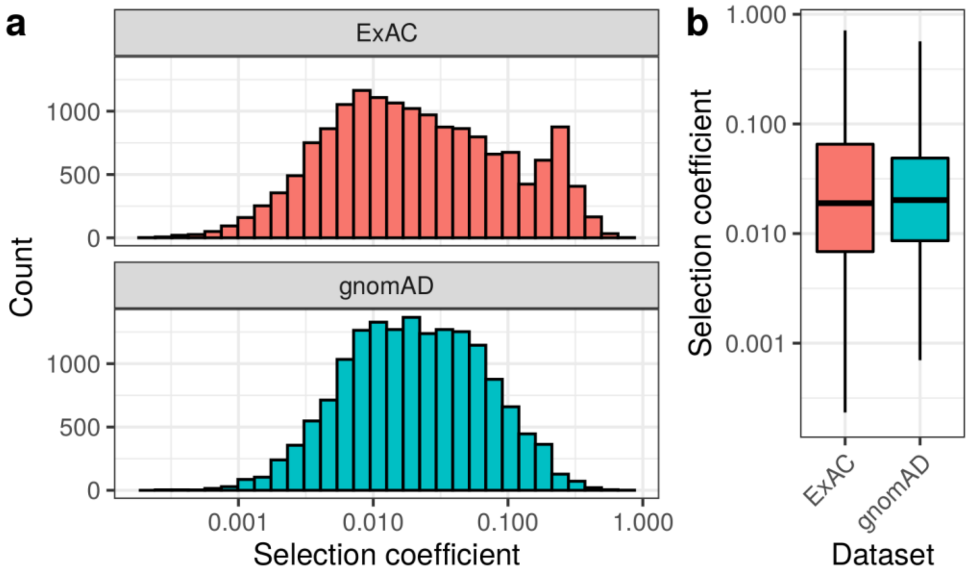


**Figure S3.** The distribution of estimated $s_{het}$ values across 15,844 genes obtained using ExAC (Cassa et al., 2017) and gnomAD data.

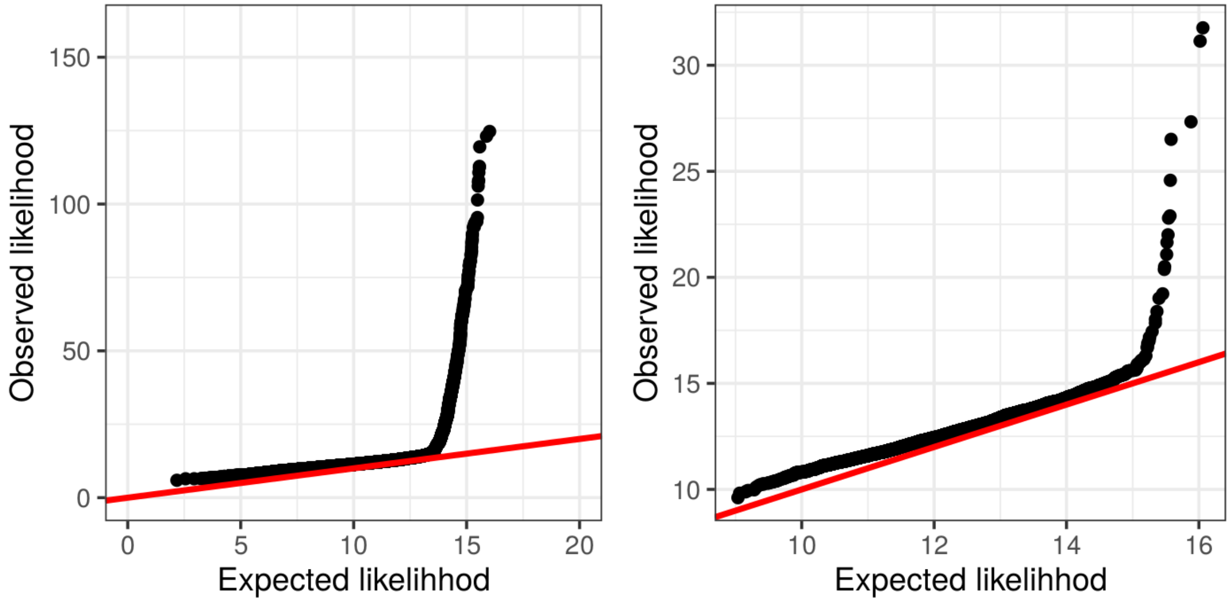


**Figure S4.** Quantile-quantile plots showing the results of the comparison between the expected and observed likelihood of a PTV count distribution. Left panel shows the results obtained for all genes, right panel - for genes with $s_{het} > 0.02$. To obtain the quantile-quantile plots, PTV count distribution across five major ancestral groups was simulated using a Poisson model, and the likelihood values were calculated for both observed and simulated data. Resulting log-likelihood scores were sorted in the ascending order, and the resulting vectors were visualized.

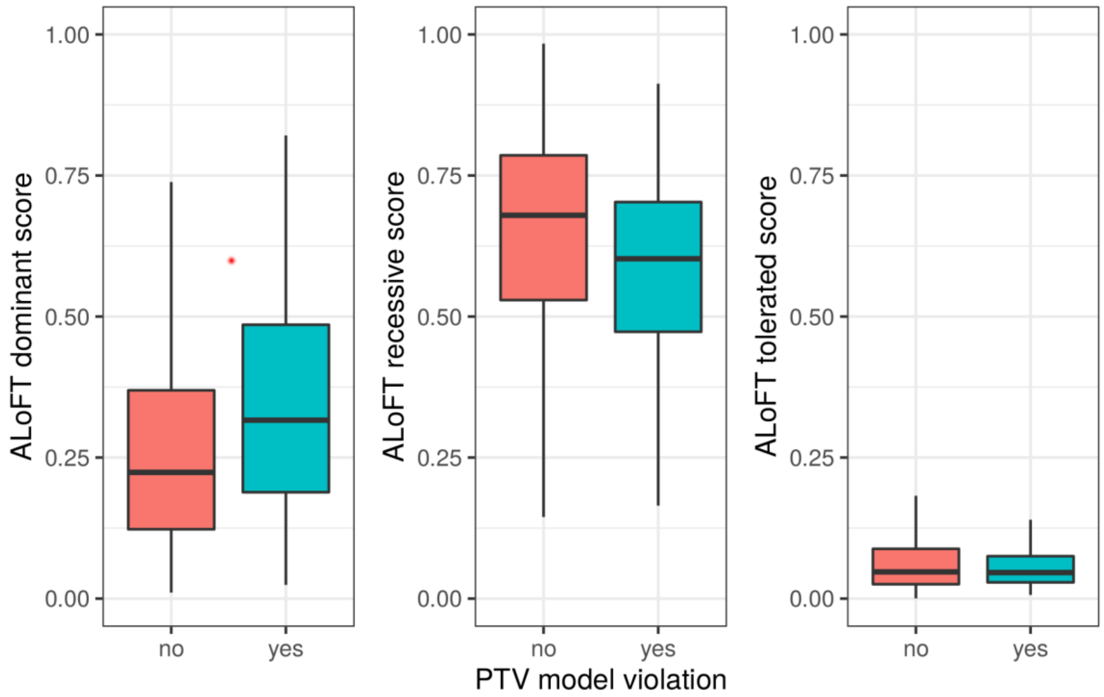


**Figure S5.** Annotation of loss-of-function transcripts (ALoFT) scores do not provide evidence of pLoF variant misclassification for genes with PTV model violation. From left to right: dominant LoF probability, recessive LoF probability, LoF-tolerated probability. The analysis was performed for genes with $s_{het} > 0.02$.

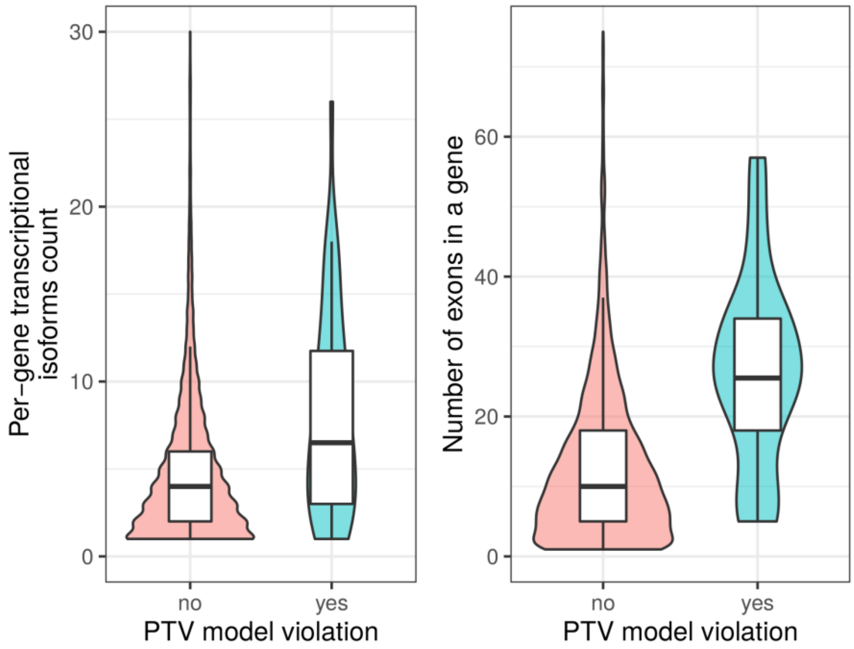


**Figure S6.** Comparison of the number of transcriptional isoforms (a) and exons (b) for "violator" and "non-violator" genes with $s_{het} > 0.02$.

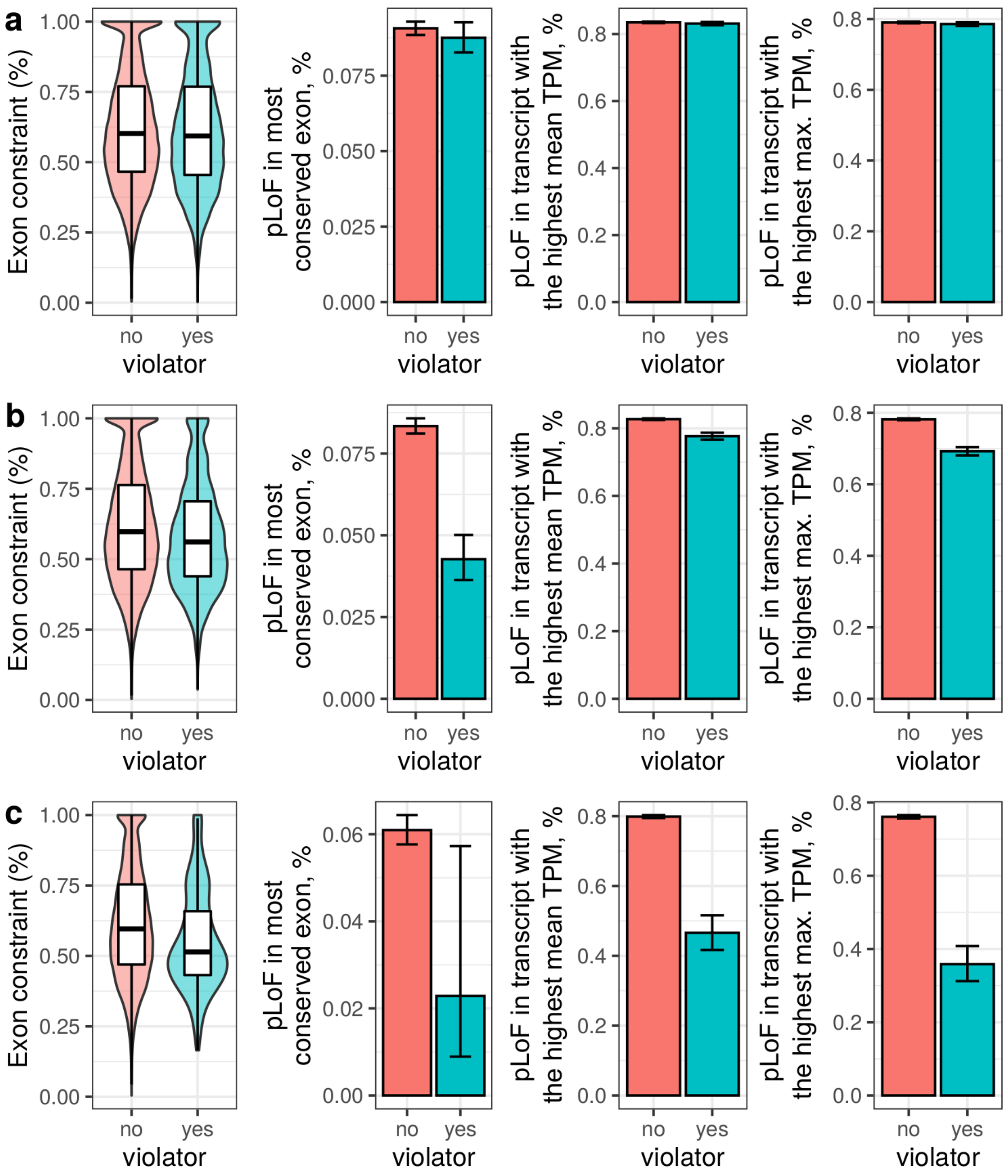


**Figure S7.** Differences between variants in "violator" and "non-violator" genes are most pronounced when considering genes with $s_{het}$ value greater than 0.02. On all panels, plots are shown in the following order (from left to right)  exon constraint as the percentage of maximum constraint within a gene; % of variants in the most constrained exon; % of variants that affect transcript with the highest average expression; % of variants affecting transcripts with the highest maximum expression. On (a), plots were generated for all genes; on (b) - for all genes with $s_{het}$ > 0.006; on (c) - for genes with $s_{het}$ > 0.02.

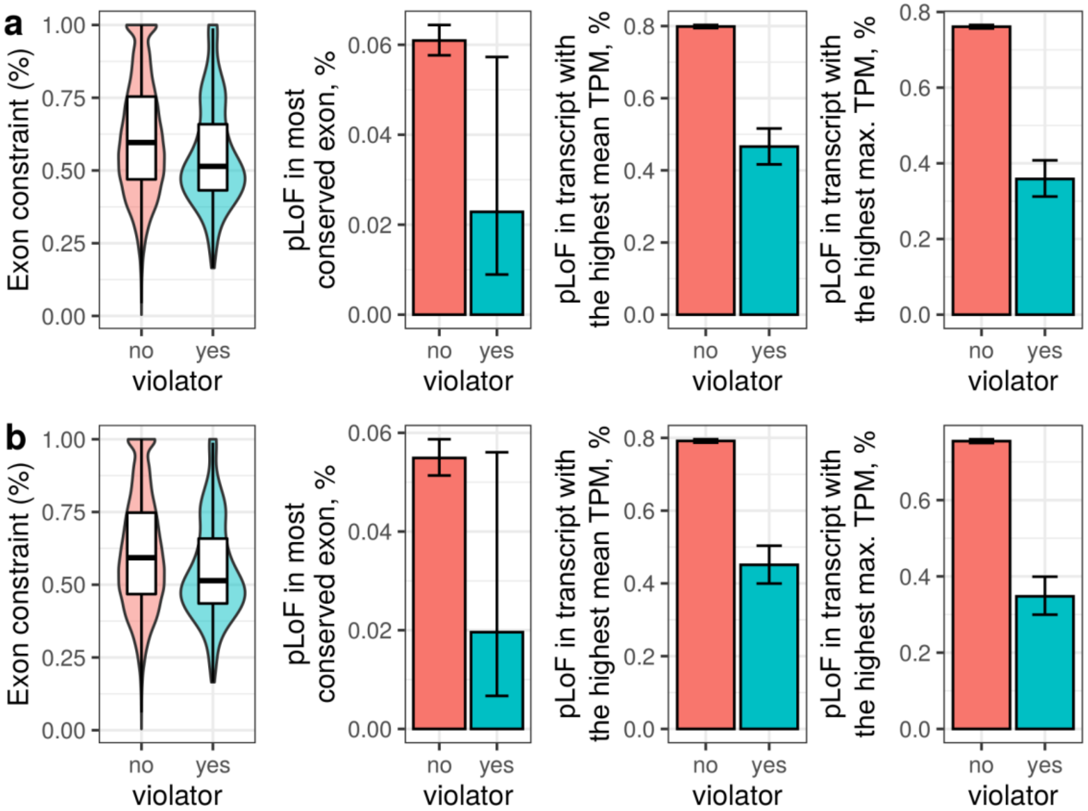


**Figure S8.** Removal of the most common PTV in each gene does not affect the functional differences between variants in "violator" and "non-violator" genes. Plots are shown in the following order (from left to right) exon constraint as the percentage of maximum constraint within a gene; % of variants in the most constrained exon; % of variants that affect transcript with the highest average expression; % of variants affecting transcripts with the highest maximum expression. (a) All variants included. (b) Most common variant in each gene removed.

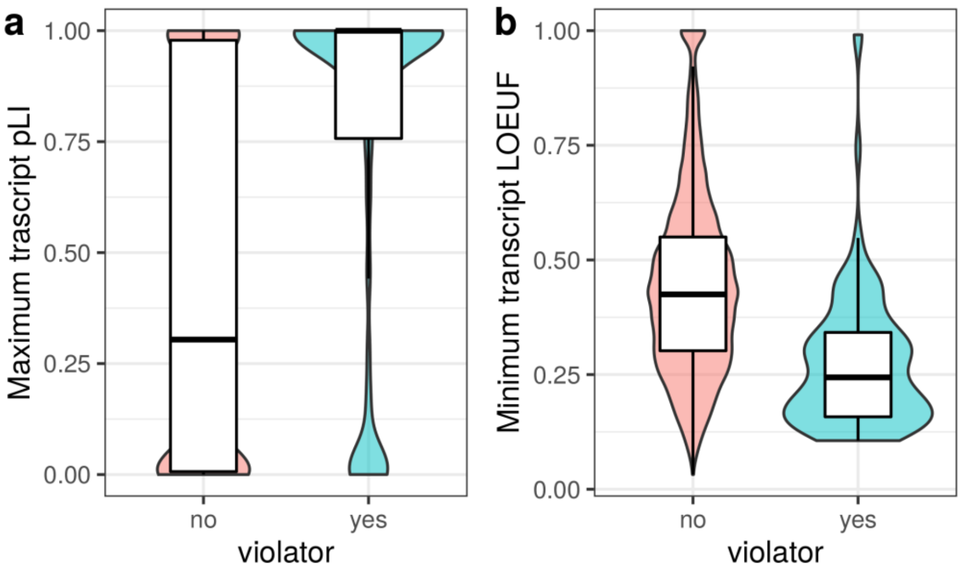


**Figure S9.** Constraint of transcripts affected by gnomAD PTVs. Shown are violin plots representing the distribution of maximum pLI (a) and minimum LOEUF (b) scores of affected transcripts for each PTV.

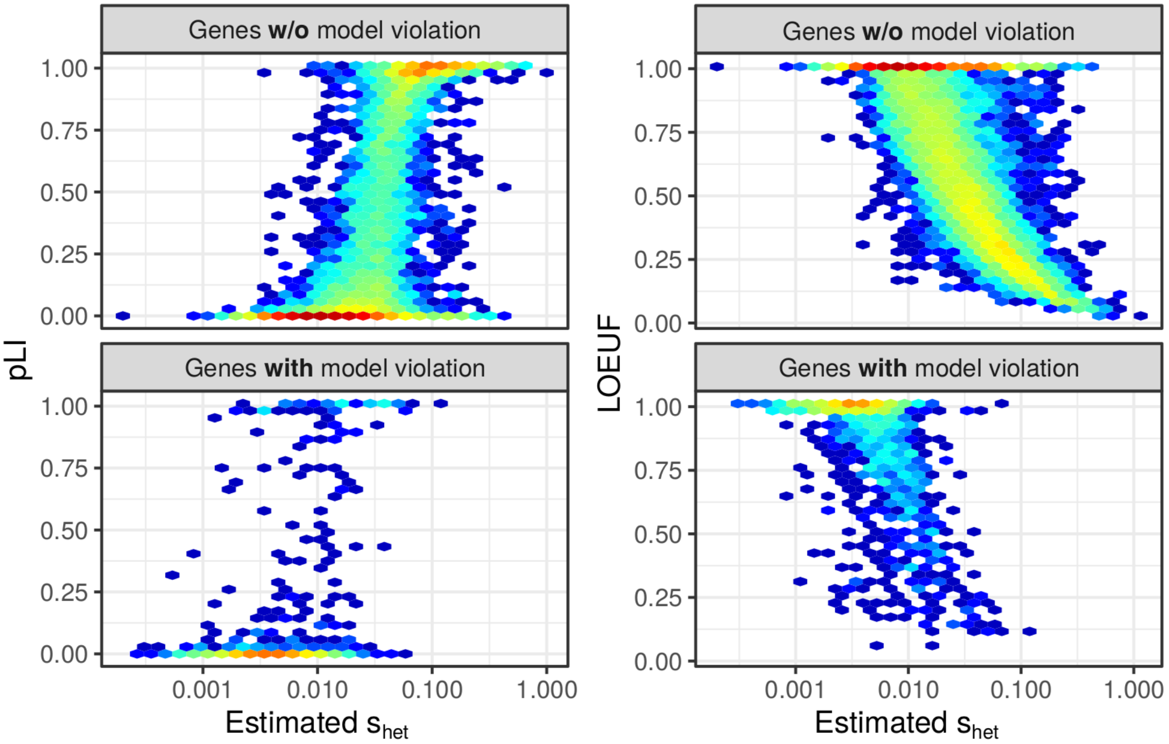


**Figure S10.** Comparison of the relationship between pLI and $s_{het}$ (left) or LOEUF and $s_{het}$ (right) for "violator" and "non-violator" genes. Hexagon color represents the $\log_{10}$ of the number of variants in the corresponding bin.

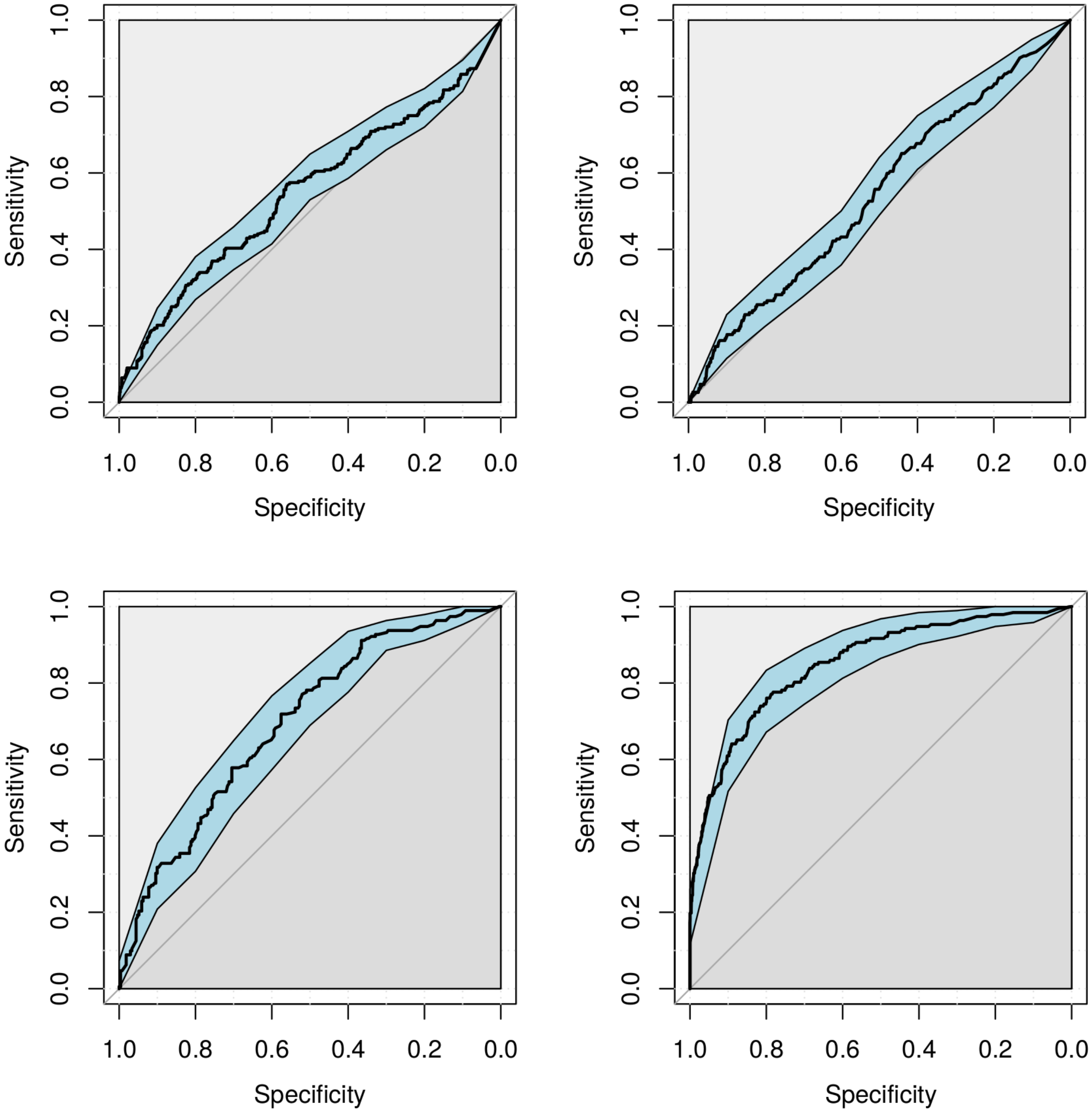


**Figure S11.** Receiver-operator curves for classification of ClinVar pathogenic vs benign variants. Top legt, pext-only model; top right, GN-C model (trained to distinguish variants in "violator" and "non-violator" genes with $s_{het}$ > 0.02); bottom left - GN-A model (similar to GN-C, but trained using all genes); bottom right, CLV model (trained using ClinVar data). ROC curves were calculated using a test subset of ClinVar variants.

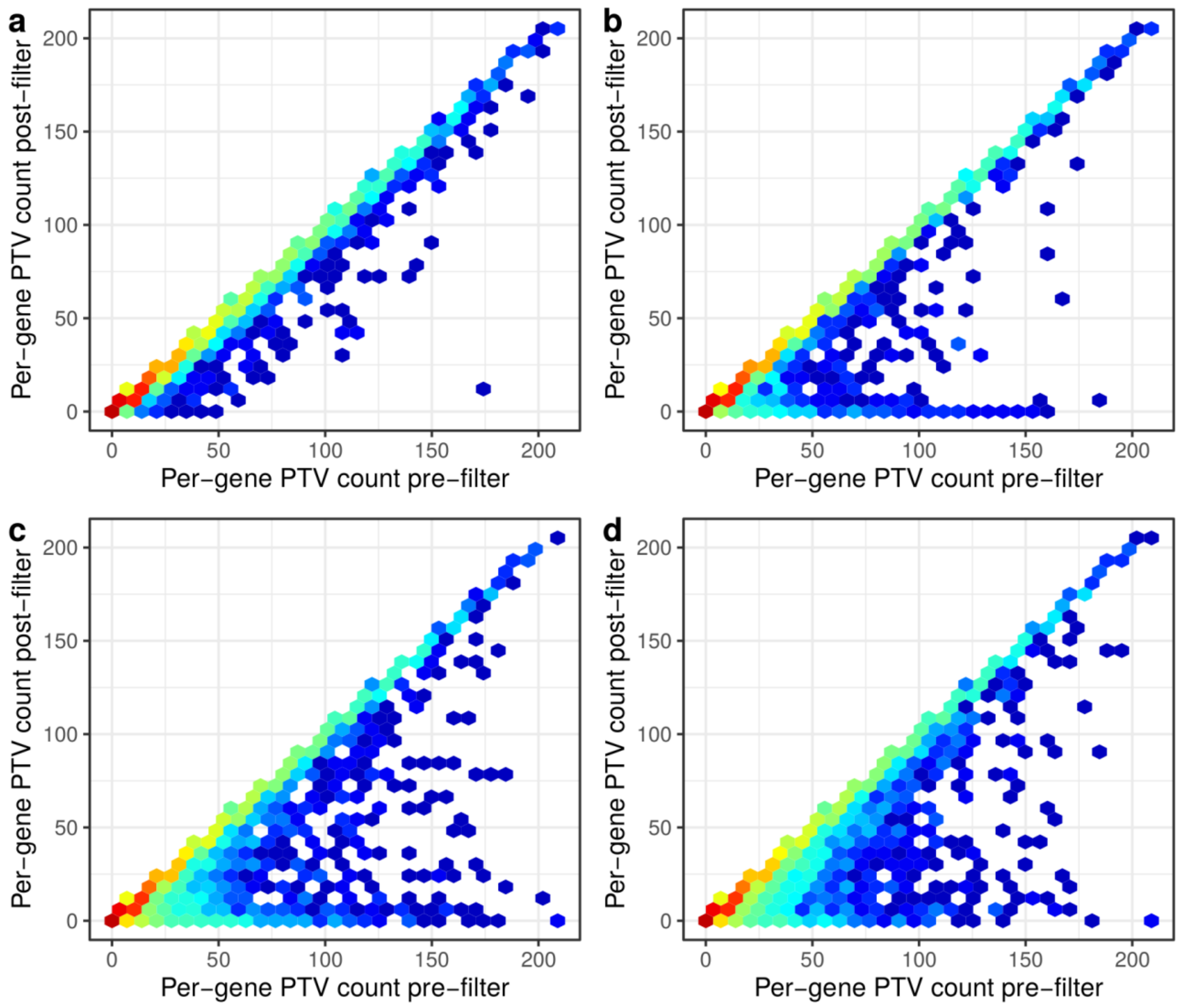


**Figure S12.** The correspondence between gene-level PTV allele counts before and after filtering of variants with various approaches: (a) random exclusion of 6,000 (~4%) variants; (b) filtering with pext < 0.1; (c) filtering with LoFfeR, GN-A model; (d) filtering with LoFfeR, CLV model. Hexagon color represents the $\log_{10}$ of the number of genes in the corresponding bin.

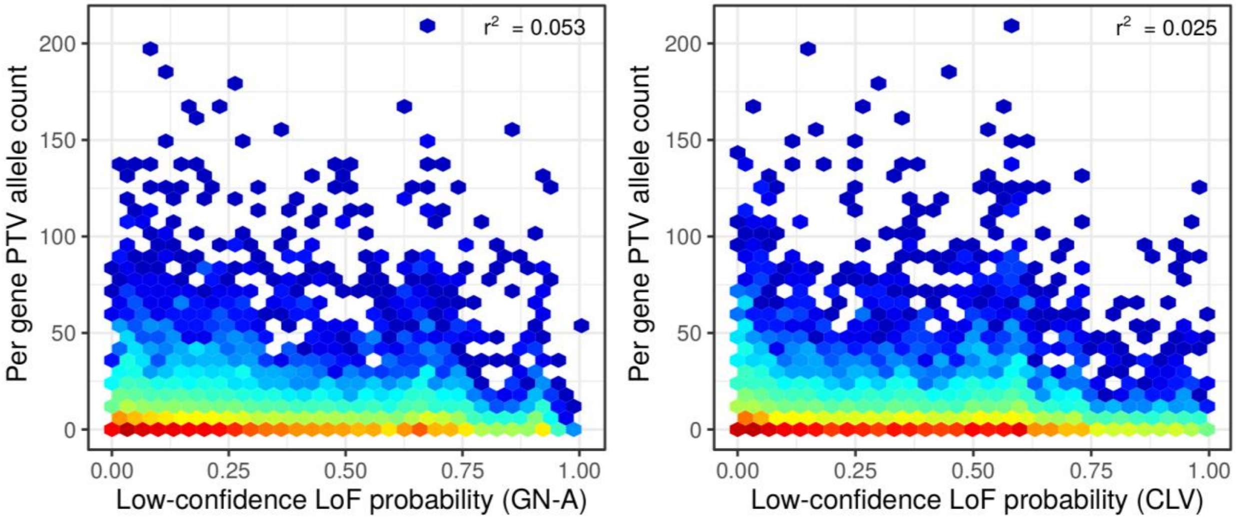


**Figure S13.** Correlation between the prediction probabilities of the GN-A (left) and CLV (right) models and the gnomAD global population allele counts (AC). Hexagon color represents the $\log_{10}$ of the number of variants in the corresponding bin.
